# Rational Design of Resistance-Suppressing Phage-Antibiotic Cocktails via Receptor Tradeoffs

**DOI:** 10.64898/2026.05.12.724642

**Authors:** Xukang Shen, Zemer Gitai

## Abstract

Bacteriophages (phages) are promising antibiotic alternatives, yet their utility is limited by rapid resistance evolution. Current phage-antibiotic combination strategies rely on empirical screening for efficacy in killing, which often fails to prevent resistance. Here, we establish a mechanistic framework to rationally design phage–antibiotic cotreatments that specifically suppress resistance by leveraging evolutionary tradeoffs. We show that traditional killing efficacy metrics do not predict resistance suppression. Instead, we identify predictable evolutionary trade-offs, including species-specific targets like TolC (*Escherichia coli*) and Type IV pili (*Pseudomonas aeruginosa*). We also identify a universal vulnerability for Gram-negative bacteria, lipopolysaccharide (LPS). We find that LPS-targeting phages are ubiquitous and that phage-steered LPS perturbation creates a predictable hypersensitivity to lipophilic antibiotics. This mechanistic trade-off, governed by physicochemical rules rather than drug-specific targets, directly correlates with resistance suppression. This framework shifts combination therapy from empirical screening to the *a priori* design of evolutionarily stable cocktails.

## Introduction

The evolution of resistance to all commonly used antibiotics has led to a global antibiotic resistance crisis^1,2^. Bacteriophages (phages) are viruses that specifically target bacteria and offer a promising alternative to traditional antibiotics^3-6^. Phages are very diverse and represent the most abundant biological entities on earth, providing a rich resource for potential therapeutic applications^7^. Additionally, phages are highly specific to bacteria and thus do not threaten human safety^8^. Despite these strengths, phage therapy has yet to become widely adopted. A major challenge in phage therapy is the emergence of bacteria that become resistant to phage infection^6,9^. The most common means by which bacteria evolve resistance to phage is by mutating the surface proteins that phages use as receptors to enter the bacteria^10-13^. This evolution appears to be clinically significant as a recent analysis of 100 clinical phage therapy cases found that all the phage-resistant bacteria analyzed carried phage receptor mutations^13^.

A promising approach to address the rise of resistance to phages would be to combine phage therapy with administration of a traditional antibiotic^13-15^. Specific phage-antibiotic combinations have been shown to both increase treatment potency and reduce resistance emergence^15^. In support of this approach, bacterial eradication is 70% less likely without the concomitant use of antibiotics^13^. However, not all antibiotic-phage combinations are effective and some combinations are even antagonistic^16-18^. We currently lack the ability to predict which of the many phages one could use in the clinic will interact well with which of the dozens of clinically-used antibiotics. The most commonly-used method to date of analyzing the interactions between phages and antibiotics is the use of synograms^19^. Synograms examine how phage and antibiotic potency affect one another in the hopes of identifying pairs whose efficacy is synergistic with respect to suppressing bacterial growth. However, we argue that this focus on killing efficacy is misplaced for long-term therapy. Efficacy synergy measures the ability to clear the *current* population but fails to account for the inevitable evolution of resistance. In fact, theoretical models suggest that efficacy synergy can accelerate multi-agent resistance because resistance to a single treatment makes it easier to acquire resistance to the second treatment^20^. To prevent treatment failure, the goal must shift from empirical screening for initial efficacy to the rational design of combinations that suppress the emergence of resistance.

One approach to directly address phage resistance emergence is to leverage evolutionary tradeoffs between phage resistance and antibiotic resistance. An evolutionary tradeoff occurs when a mutation beneficial for one trait is detrimental for another^21^. For example, a previous study demonstrated that when a phage (like U136B) targets an antibiotic efflux (like TolC), then mutants that become resistant to the phage by losing the efflux pump become more sensitive to the antibiotics that are effluxed by that pump^22^. Unfortunately, relatively few phages use efflux pumps as receptors and efficiently-effluxed antibiotics are less favored clinically, such that the utility of this specific example has remained limited. It has thus remained unclear whether resistance tradeoffs can be extended to other phages and if they can be used to predict effective outcomes in phage–antibiotic cotreatments.

Here, we sought to develop a rational design framework for predicting resistance-suppressing phage-antibiotic combinations based on evolutionary tradeoffs. We systematically characterized how resistance to phylogenetically diverse phages alters *E. coli* antibiotic susceptibility using receptor knockout mutants and experimental evolution. We found that very few *E. coli* phages target efflux pumps, but that the majority of *E. coli* phages do target receptors whose loss leads to predictable changes in antibiotic sensitivity. Specifically, the outer membrane lipopolysaccharide (LPS) represents a key vulnerability because more than half of all *E. coli* phages target LPS and phages steer the evolution of LPS mutants that sensitize those bacteria to lipophilic antibiotics that otherwise struggle to traverse the outer membrane. Moreover, we found that the evolutionary tradeoffs we identified predictably suppress the emergence of resistance during phage–antibiotic cotreatments. Finally, we confirmed the generality of this approach by identifying genetic tradeoffs between a different set of phages and antibiotics targeting *Pseudomonas aeruginosa* and confirming that these tradeoffs effectively predict resistance suppression in this clinically important pathogen.

## Results

### Deletion of Phage Receptors Alters Antibiotic Susceptibility in Escherichia coli

Because most phage resistance mutations are predictably loss-of-function mutations in phage receptors, we sought to understand how deletion of these receptors affects bacterial sensitivity to various antibiotics. We examined the BASEL collection of diverse *E. coli* phages that represents the natural diversity of *E. coli* phages^23^. In the original study describing this collection, receptor identification was performed using spotting assays on a set of *E. coli* strains lacking specific surface-exposed outer membrane factors. To validate these receptor assignments, we isolated *E. coli* mutants resistant to 14 representative phages from the major BASEL phage groups and sequenced their genomes to identify the resistant mutations (**Table S1**). Sequencing the mutations in these resistant mutants largely confirmed the original receptor annotations with one exception, in which we found that Bas1, which was originally annotated as an LPS-targeting phage, elicited mutations in the outer membrane protein, NupG (**Fig. S1**). To further confirm the original annotations of the receptor identities for all of the BASEL phages, we used a cross-streaking assay to test each phage’s ability to infect a deletion mutant in its predicted receptor. Across the 62 phages in the BASEL collection, the majority (33, 53.2%) use lipopolysaccharide (LPS) as a receptor, while the remainder use outer membrane proteins, including BtuB (11, 17.7%), FhuA (10, 16.1%), LamB (4, 6.5%), YncD (2, 3.2 %), TolC (1, 1.6%), and NupG (1, 1.6%) (**Fig. 1a**). We note that these findings reinforce the conclusion that phages that target efflux pump receptors (like TolC) are rare (**Fig. 1a**).

**Fig. 1.**
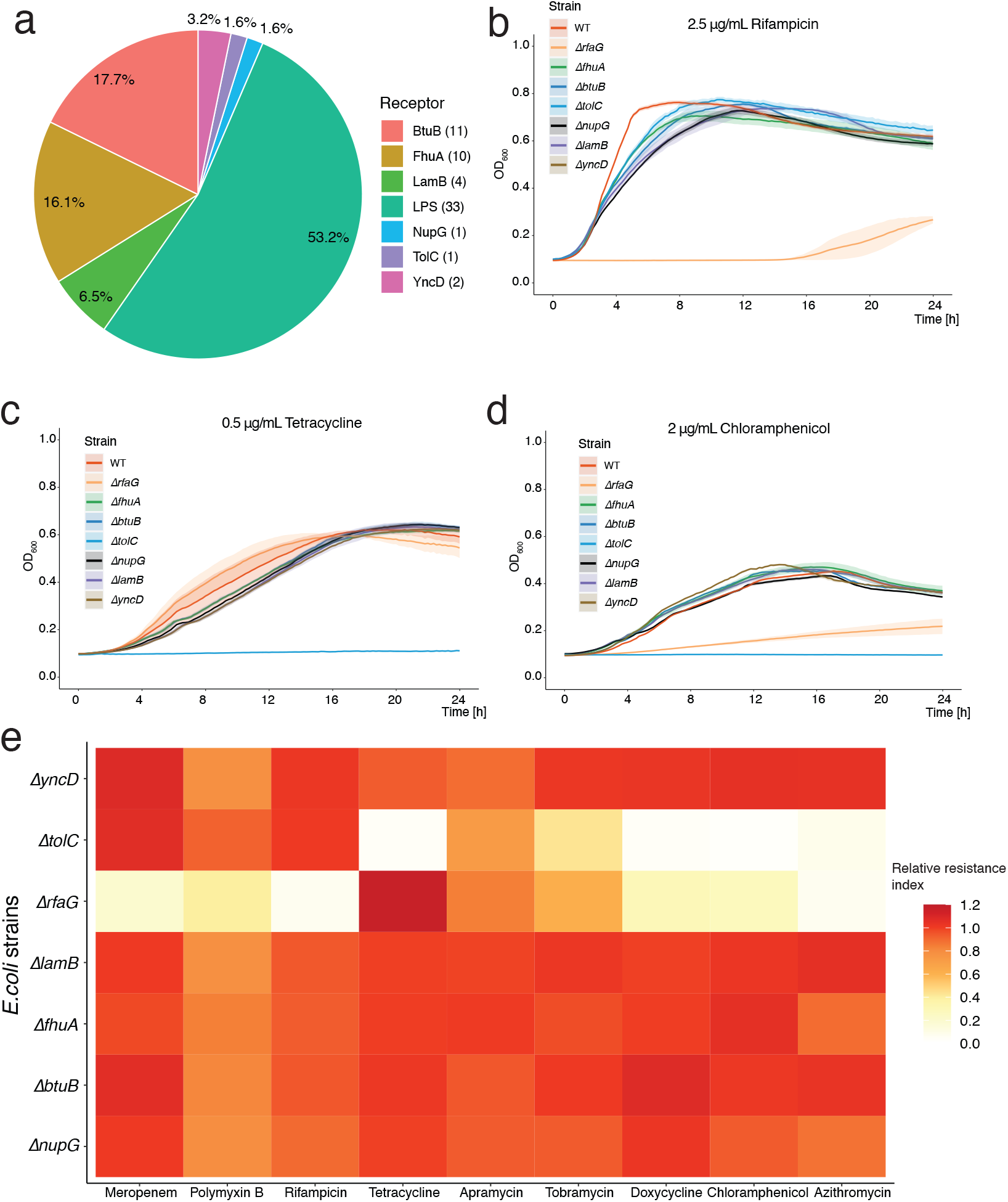
Phage receptor deletion changes *E. coli* antibiotic susceptibility. **a**, The distribution of BASEL phage receptors, showing their counts in parentheses and percentages in the pie chart. **b, c, d**, Growth curves of receptor-knockout mutants and wild-type *E. coli* in one fourth of the MIC for rifampicin (**b**, 2.5 µg/mL), tetracycline (**c**, 0.5 µg/mL), and chloramphenicol (**d**, 2 µg/mL). Curves represent the mean of three independent biological replicates ± s.e.m. **e**, Relative resistance index of phage receptor mutants compared to wild-type *E. coli* across nine antibiotics. In the heat map, each column represents an antibiotic and each row shows a phage receptor deletion mutant strain. Color indicates the mean of relative resistance index values of three biological replicates.

Because the phage resistant mutants were largely loss-of-function mutants, we sought to systematically assess the impact of phage receptor deletion on antibiotic susceptibility. We used Δ*rfaG* to assay LPS deficiency, as this strain was previously shown to resist infection by BASEL phages that use LPS as their receptors^23^. We tested the sensitivity of these phage receptor deletion mutants and wild-type *E*.*coli* against a panel of nine antibiotics, each at one fourth of the wild-type minimum inhibitory concentration (MIC). These antibiotics spanned diverse targets and chemical properties: transcription (rifampicin)^24^, translation (azithromycin, chloramphenicol, doxycycline, tetracycline, apramycin, tobramycin)^25-27^, membrane integrity (polymyxin B)^28^, and cell wall synthesis (meropenem)^29^. We found that phage receptor deletions conferred distinct patterns of antibiotic susceptibility. Most notably, LPS deficiency (Δ*rfaG*) increased sensitivity to rifampicin (**Fig. 1b**), meropenem (**Fig. S2a**), and polymyxin B (**Fig. S2b**). The polymyxin B sensitivity is particularly notable because LPS-associated mutations have also been implicated in polymyxin B resistance^30^, demonstrating that phage-resistance leads to a distinct class of LPS alteration. Deletion of *tolC* specifically increased susceptibility to tetracycline (**Fig. 1c**) and both Δ*rfaG* and Δ*tolC* mutants exhibited modest growth defects in tobramycin (**Fig. S2c**) and apramycin (**Fig. S2d**) but showed pronounced growth inhibition in chloramphenicol (**Fig. 1d**), doxycycline (**Fig. S2e**) and azithromycin (**Fig. S2f**).

To systematically compare how each phage receptor deletion mutant affects sensitivity to the antibiotics tested, we calculated a “relative resistance” index by calculating the ratio of the area under the growth curve (AUC) for each mutant relative to wild type for each antibiotic at one quarter (0.25X) of the wild type MIC of that antibiotic. For each antibiotic, mutants that do not affect antibiotic resistance would have a relative resistance index of 1.0, mutants with increased sensitivity have relative resistance indexes below 1, and mutants with increased resistance have relative resistance indexes above 1. The LPS-perturbed Δ*rfaG* mutant became sensitive to most antibiotics tested, with the notable exception of tetracycline, which showed increased relative resistance (**Fig. 1e**). Strong sensitivity (relative resistance <0.5) was observed for meropenem, polymyxin B, rifampicin, doxycycline, chloramphenicol, and azithromycin. Meanwhile, Δ*tolC* strongly sensitized *E. coli* to tetracycline, tobramycin, doxycline, chloramphenicol, and azithromycin. For the remaining receptor mutants (Δ*btuB*, Δ*fhuA*, Δ*lamB*, Δ*yncD*, Δ*nupG*), relative resistance was above 0.5 for all antibiotics tested, suggesting minimal changes in antibiotic susceptibility (**Fig. 1e**).

### LPS mutant antibiotic sensitivity correlates with antibiotic lipophilicity

The resistance patterns of Δ*rfaG* and Δ*tolC* were qualitatively distinct with respect to multiple antibiotics, suggesting that they affect antibiotic sensitivity through distinct mechanisms. TolC is a well-characterized efflux pump and our findings were largely consistent with TolC’s known effects on antibiotic sensitivity^31^. The effect of perturbing LPS on antibiotic resistance is less clear because LPS can act both as a direct barrier to antibiotic entry^32^ and as an indirect mediator of the assembly of outer membrane proteins^33^. We thus sought to determine if the patterns of antibiotic sensitization we observed could be explained by the biochemical properties of the antibiotic or their mechanisms of action. Doxycycline (C_22_H_24_N_2_O_8_) is a semi-synthetic derivative of tetracycline (C_22_H_24_N_2_O_8_)^34^. Both antibiotics inhibit protein translation by binding to the 30S ribosomal subunit^26^. However, LPS deficiency renders *E. coli* more susceptible to doxycycline but not to tetracycline (**Fig. 1e**). Structurally, doxycycline lacks the C-6 hydroxyl group found in tetracycline and instead has a hydroxyl group shifted to C-5^35^. This modification increases its lipophilicity (doxycycline logP = –0.46 vs. tetracycline logP = –0.56)^36^. LPS has been proposed to act as a barrier to the cell entry of lipophilic molecules, which would explain why the LPS defects would preferentially enhance the entry of doxycycline. Consistent with this, we observed a significant negative correlation between antibiotic lipophilicity and the relative resistance indexes of the *ΔrfaG* mutant (ρ = -0.90, *P* = 0.005), indicating that LPS deficiency sensitizes cells preferentially to lipophilic antibiotics (**Fig. S3**).

### Experimentally evolved phage resistance predictably alters antibiotic sensitivity

Because deletions of phage receptors can alter antibiotic susceptibility, we next investigated whether bacteria that evolve phage resistance also exhibit similar changes in antibiotic susceptibility. To test this, we performed experimental evolution of *E. coli* with seven distinct *E. coli* phages, each targeting a different receptor, representing the complete set of phage receptors in the BASEL collection. We performed three days of serial passaging because all isolated *E*.*coli* colonies from day-three populations were resistant to their respective coevolving phages, indicating that phage resistance had evolved (**Fig. 2a**). Phages could still be detected in all populations, indicating ongoing selection pressure from phages. We note that all the phages in the BASEL collection are lytic such that we did not need to account for lysogeny (and no lysogeny was observed).

**Fig. 2.**
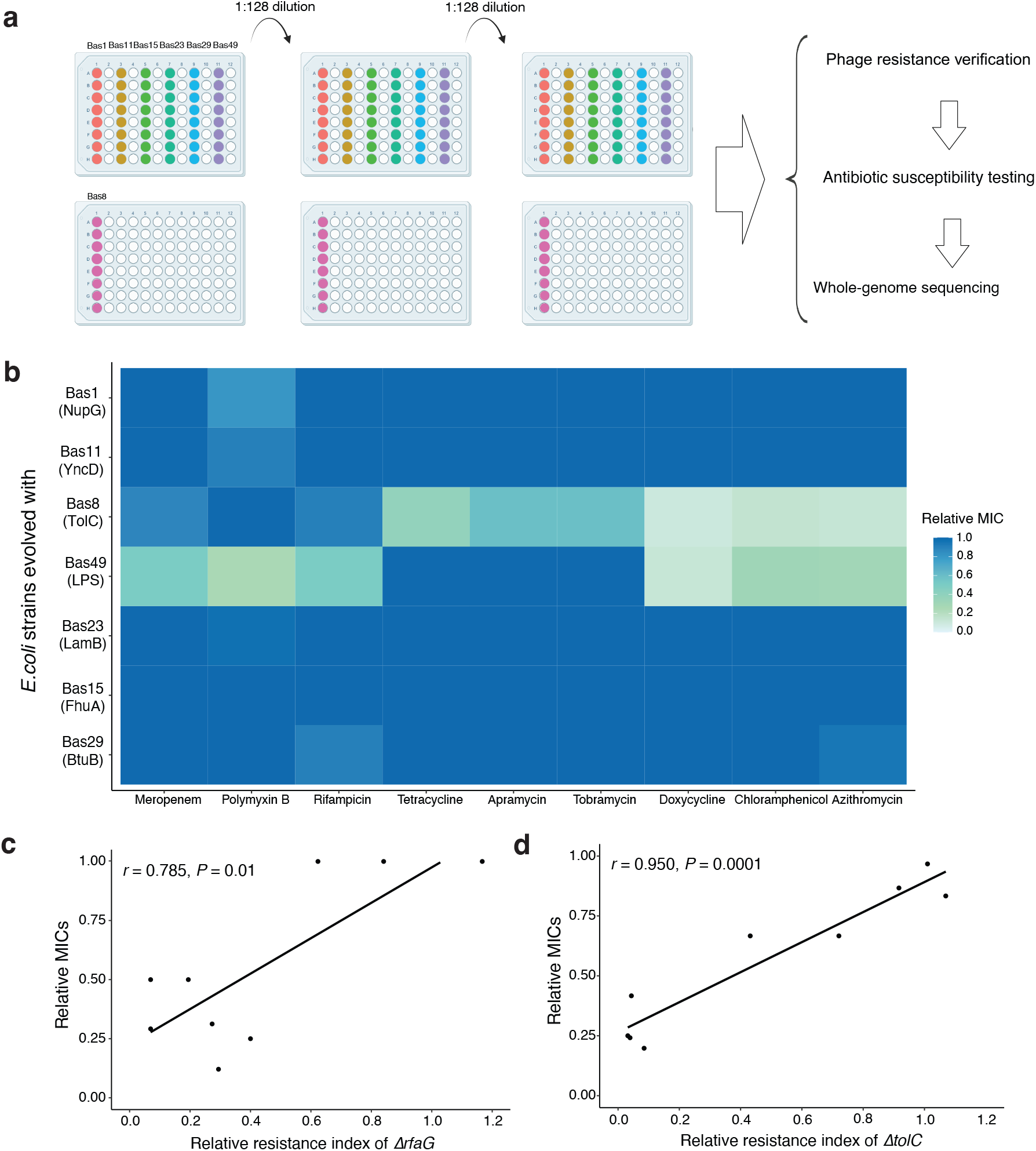
Phage resistance shifts antibiotic sensitivity. **a**, Schematic of *E. coli* experimental evolution with seven BASEL phages targeting different receptors, followed by phenotypic and genetic analyses of evolved clones. **b**, Relative MICs of *E. coli* evolved with seven phages across nine antibiotics. Colors represent the mean relative MIC of twelve clones (two clones from each of six independent evolution populations). **c**, Positive correlation between relative resistance index of *ΔrfaG* and relative MICs of *E. coli* evolved with phage Bas49 for nine antibiotics. **d**, Positive correlation between relative resistance index of *ΔtolC* and relative MICs of *E. coli* evolved with phage Bas8 for nine antibiotics.

To uncover the genetic basis of resistance, we performed whole-genome sequencing on multiple evolved *E. coli* clones isolated from each evolution experiment. Of the 43 isolates sequenced, 37 carried mutations in the expected phage receptor and 5 carried mutations in known transcriptional regulators of the expected phage receptor (*i*.*e. malT* is a known regulator of *lamB*^37^) (**Table S2**). The only strain without a mutation in its expected receptor or receptor regulator was a single isolate of a strain resistant to Bas01, which carried a deletion in *dcrB*, a gene known to be involved in phage DNA injection^38^. Interestingly, none of the 43 isolates carried simple point mutations: 17 displayed deletion mutations, 18 displayed transposon mutations, 5 displayed premature stop mutations, and 3 displayed frameshift mutations (**Table S2**). These results indicate that the vast majority of phage resistance is driven by mutations in phage receptors and that these mutations are largely significant genomic alterations that are likely to lead to strong loss of function of the receptor structures.

Because the mutants in the evolved phage-resistant strains were primarily loss-of-function mutations, we expected their phenotypes to resemble those of deletion mutants in the corresponding receptors. To assess this prediction, we assayed the antibiotic sensitivity of our experimentally evolved phage resistant mutants. We isolated two independent colonies from the day-three phage-resistant populations for each phage and measured the MICs of each evolved clone against the nine antibiotics previously tested on phage receptor deletion mutants. Relative MICs were calculated to compare sensitivity of the evolved strains to the wild-type progenitor strain for each antibiotic (**Fig. 2c)**. The changes in MIC patterns of these evolved phage-resistant mutants closely mirrored the relative resistance index patterns observed for the receptor deletion mutants. For example, correlation analyses confirmed that the antibiotic susceptibility patterns of Bas8- and Bas49-resistant clones were significantly correlated with those of the corresponding receptor mutants (*p* < 0.05, **Fig. 2d** and **2e**).

More than half of all BASEL phages use LPS as a receptor and these LPS-targeting phages are phylogenetically diverse. To determine whether the antibiotic susceptibility profiles of *E*.*coli* resistant to Bas49 were representative of *E*.*coli* resistant to LPS-targeting phages more broadly, we evolved *E. coli* with eight additional LPS-targeting phages that belong to two major phage groups (Siphoviridae: Bas19; and Myoviridae: Bas46, Bas48, Bas50, Bas51, Bas52, Bas53 and Bas54) and measured MICs of resistant clones. In each case, the evolved strains exhibited similar antibiotic susceptibility patterns to those of Bas49-resistant cells (**Fig. S4**).

Thus, while LPS-targeting phages are phylogenetically diverse, they all show the same predictable patterns of antibiotic sensitization. Furthermore, antibiotic lipophilicity, which predicted the relative resistance indexes of antibiotics in the Δ*rfaG* strain (**Fig. S3**), also predicted antibiotic MICs in *E. coli* that evolved resistance to these diverse LPS-targeting phage (**Fig. S5**).

### Evolutionary tradeoffs predict resistance suppression in phage-antibiotic cotreatments

Can the tradeoffs between phage resistance and antibiotic sensitivity be leveraged to predict phage–antibiotic cotreatments that suppress resistance emergence? To address this question, we examined whether phage–antibiotic combinations exhibiting strong tradeoffs also showed reduced resistance during cotreatment. We cultured wild-type *E. coli* (5 × 10^6^ cells/mL) with different phages (10^6^ phages/mL) and antibiotics at subinhibitory concentrations (0.25 × wild-type MIC) (**Fig. 3a-b** and **Fig. S6**). When grown with phage alone, bacterial populations displayed a reproducible pattern with three phases: an initial rise in density (reflecting bacterial growth initially exceeding phage infection), followed by a sharp collapse (driven by phage replication and lysis), and finally, a regrowth phase. Cross-streak assays confirmed that this late regrowth phase represented the emergence of phage-resistant cells (**Fig. S7**).

**Fig. 3.**
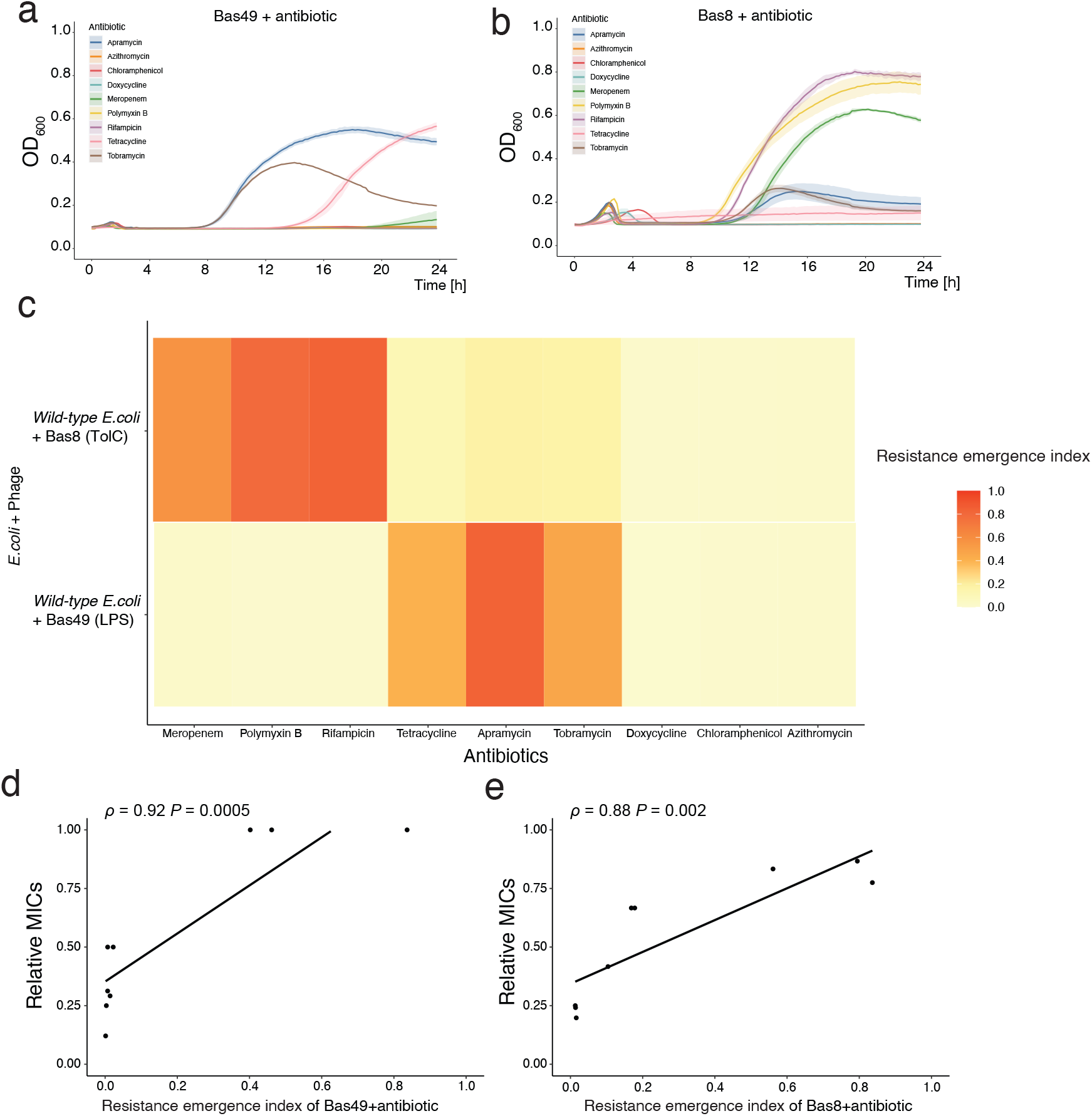
Tradeoffs predict resistance emergence in cotreatments. **a**, Growth curves of *E. coli* cotreated with phage Bas49 and nine antibiotics. Curves represent the mean of three independent biological replicates ± s.e.m. **b**, Growth curves of *E. coli* cotreated with phage Bas8 and nine antibiotics. Curves represent the mean of three independent biological replicates ± s.e.m. **c**, Relative resistance indexes of Bas49, antibiotic cotreatments and Bas8, antibiotic cotreatments. Colors indicate relative resistance index values from three biological replicates. **d**, Positive correlation between resistance emergence indexes of Bas49–antibiotic cotreatments and relative MICs of *E. coli* evolved with phage Bas49 across the nine antibiotics. **e**, Positive correlation between resistance emergence indexes of Bas8–antibiotic cotreatments and relative MICs of *E*.*coli* evolved with phage Bas8 across the nine antibiotics.

For the phages targeting receptors whose mutants did not exhibit significant antibiotic sensitivity, we did not expect to observe significant suppression of resistance emergence. As predicted, in the presence of antibiotics, resistance robustly emerged in all combinations involving phages tested targeting receptors other than LPS and TolC: Bas11 (targeting YncD), Bas15 (FhuA), Bas23 (LamB), and Bas29 (BtuB) (**Fig. S6**). In contrast, antibiotic combinations involving the TolC-targeting phage Bas8 or LPS-targeting Bas49 showed variable outcomes—resistance appeared in some antibiotic pairings but was largely suppressed in others. To quantify resistance emergence in these cotreatments, we calculated a *resistance emergence index* as the AUC of the late resistance-associated growth peak found in the presence of a given antibiotic, normalized to the AUC of the resistance-associated peak observed in phage-only conditions (**Fig. S8**). High resistance emergence indexes (approaching 1.0) indicate that a given phage-antibiotic combination does not suppress the emergence of resistance while low resistance indexes (approaching 0.0) indicate strong resistance suppression. For Bas49 cotreatments, resistance emergence indexes were high with tetracycline, apramycin and tobramycin, but much lower with meropenem, polymyxin B, rifampicin, doxycycline, chloramphenicol, and azithromycin **(Fig. 3c)**. In contrast, Bas8 cotreatments showed high indexes with meropenem, polymyxin B, and rifampicin, but low values with tetracycline, apramycin, tobramycin, doxycycline, chloramphenicol, and azithromycin. These results indicate that with respect to suppressing the emergence of resistance, both Bas8 and Bas49 synergize well with doxycycline, chloramphenicol, and azithromycin, while Bas49 specifically synergizes well with meropenem, polymyxin B, and rifampicin, and Bas8 specifically synergizes well with tetracycline, apramycin, and tobramycin **(Fig. 3c)**.

The specificity observed with respect to suppressing resistance emergence, in which different antibiotics synergize differently with different phages, highlights the current challenge in constructing effective phage-antibiotic cocktails. Phage-antibiotic combinations are traditionally assessed by the extent of initial bacterial eradication rather than focusing on resistance emergence. We thus asked whether resistance tradeoffs or initial cotreatment efficacy better correlate with resistance emergence. We found that resistance emergence indexes were significantly negatively correlated with the MICs of phage-resistant strains for both Bas49–antibiotic combinations (**Fig. 3d**) and Bas8–antibiotic combinations (**Fig. 3e**) cotreatments. The ability of Bas49-antibiotic combinations to suppress resistance emergence also correlated with the lipophilicity of the antibiotic used **(Fig. S9)**, supporting our hypothesis that this occurs because mutations that perturb LPS to reduce phage entry simultaneously make it easier for lipophilic antibiotics to enter the bacteria.

In our combination treatment growth curves, initial cotreatment efficacy is represented by the AUC for the initial growth peak in the phage-antibiotic combination treatment relative to the corresponding initial peak in the phage treatment alone. This cotreatment efficacy relative AUC did not significantly correlate with the resistance emergence indexes for Bas49 cotreatments (ρ = 0.27, *P* = 0.49) or Bas8 cotreatments (ρ = -0.42, *P* = 0.27) (**Fig. S10**). Since LPS-targeting phages represent the majority of the phages in the BASEL collection and exhibit strong antibiotic tradeoffs, our results suggest that for many *E. coli* phages antibiotic resistance tradeoffs could be used to predict phage-antibiotic cocktails that suppress resistance emergence more reliably than cotreatment efficacy.

### Phage-antibiotic tradeoffs also predict resistance emergence for Pseudomonas aeruginosa

Having established a strategy to identify effective phage–antibiotic cotreatments that suppress resistance emergence in *E. coli* through resistance tradeoffs, we next asked whether this approach could be generalized to a clinically important pathogen, *Pseudomonas aeruginosa. P. aeruginosa* is an opportunistic pathogen that can cause infections in the blood, lungs (pneumonia), urinary tract, and other sites^39^. In 2017, multidrug-resistant (MDR) *P. aeruginosa* was responsible for an estimated 32,600 infections among hospitalized patients and 2,700 deaths in the United States^40^. *P. aeruginosa* infections are often chronic^41^, such that it is feasible to culture isolates from patients and develop patient-specific treatments. In a systematic analysis of receptors used by 27 *P. aeruginosa* phages collected from sewage, 23 used LPS as their receptors and only 4 were found to target type IV pili (T4P)^42^ (**Fig. 4a**). This distribution indicates that like *E. coli*, LPS represents a major target for *P. aeruginosa* phages.

**Fig. 4.**
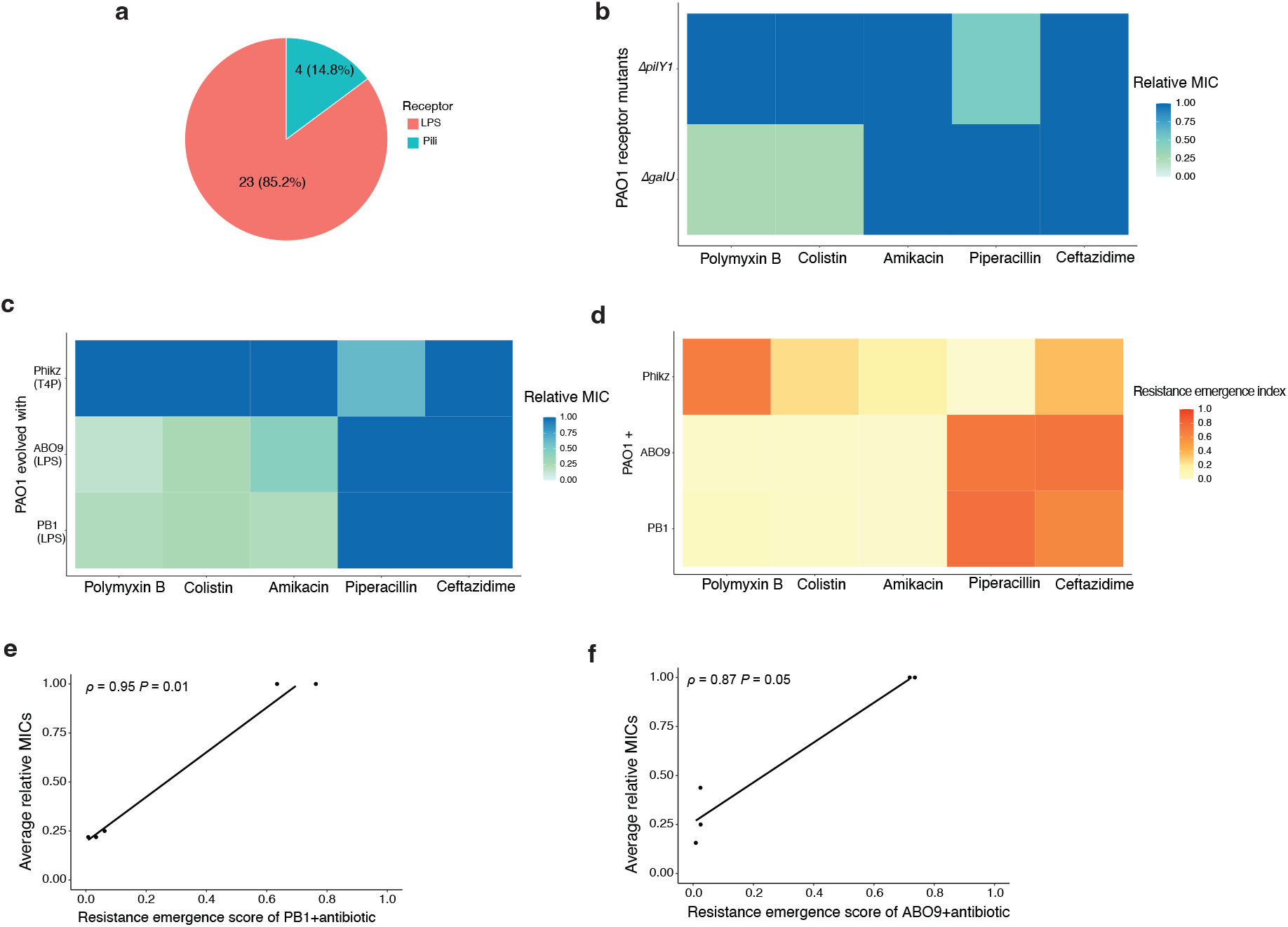
Applying the tradeoff-based strategy to *Pseudomonas aeruginosa*. **a**, Receptor usage of 27 *P. aeruginosa* phages isolated from natural environments (data from Wright *et al*., 2018). **b**, Relative MICs of phage receptor mutants (Δ*galU*, Δ*pilY1*). In the heat map, each column represents an antibiotic and each row represents a mutant strain. Colors indicate the average relative MICs from three biological replicates. **c**, Relative MICs of PAO1 strains evolved to be resistant to three phages across five antibiotics. Colors represent the average of four resistant clones. **d**, Relative resistance emergence indexes of PhiKZ, PB1, ABO9 and antibiotic cotreatments. Colors indicate relative resistance indexes from three biological replicates. **e**, Positive correlation between relative resistance index of PB1–antibiotic cotreatments and relative MICs of PAO1 strains resistant to phage PB1 across five antibiotics. **f**, Positive correlation between relative resistance indexes of ABO9-antibiotic cotreatments and relative MICs of PAO1 strains resistant to phage ABO9 across five antibiotics.

Since LPS and T4P represent the primary receptors for *P. aeruginosa* phages, we sought to determine if mutants that perturb LPS or T4P alter *P. aeruginosa* antibiotic sensitivity. We generated *P. aeruginosa* PAO1 deletion strains in *galU* to disrupt LPS^43^ and *pilY1* to disrupt T4P^44,45^. We confirmed that *ΔgalU* conferred resistance to the LPS-targeting phages, ABO9 and PB1 and that *ΔpilY1* conferred resistance to the T4P-targeting phage, PhiKZ. We then tested the antibiotic susceptibility of these mutants against five antibiotics commonly used in *P. aeruginosa* treatment: polymyxin B, colistin, amikacin, piperacillin, and ceftazidime^46^ (**Fig. 4b**). Deletion of *galU* increased susceptibility to B and colistin while *pilY1* increased susceptibility to piperacillin (**Fig 4b)**.

We next determined if the antibiotic sensitivities of *ΔgalU* and *ΔpilY1* are representative of the sensitivities that arise upon the evolution of phage resistance. We experimentally evolved 4 independent populations with each of the three phages ABO9, PB1, and PhiKZ, serially passaged the cultures for 3 days, isolated a single phage-resistant colony for each independent culture, confirmed that isolate to be phage-resistant, and tested its sensitivity to the five antibiotics examined with the phage receptor deletion mutants. All of the PB1-resistant (n = 4) and ABO9-resistant (n = 4) strains exhibited increased sensitivity to colistin and polymyxin B, as was expected from the *ΔgalU* sensitivity. However, unlike the case for *ΔgalU*, many (n = 7) of these mutants were also sensitized for amikacin **(Table S3** and **Fig. 4c**). Whole-genome sequencing revealed that all of the amikacin-sensitive strains carried large genomic deletions that deleted *galU* in addition to neighboring genes including the known antibiotic transporter genes *mexXY*^47^. To determine if the loss of *mexXY* might explain the amikacin sensitivity we constructed a Δ*mexX*Δ*mexY* double-deletion strain and found it to be more sensitive to amikacin (**Fig. S11**). Consistent with the hypothesis that *galU* mutants sensitize PAO1 to colistin and polymyxin B but not amikacin, the one ABO9 phage-resistant mutant we isolated that was not amikacin-sensitive harbored a point mutant in *galU* with no genetic changes in *mexXY* **(Table S3)**. All of the PhiKZ-resistant strains (n = 4) carried mutations in T4P genes (*pilI, pilJ*, or *pilY1*) or T4P regulators (*retS*) **(Table S3)**. The three T4P gene mutants displayed strong sensitization to piperacillin, similar to that of *ΔpilY1* **(Fig. 4c and Table S3)**. The *retS* mutant did not show significant sensitization to any of the antibiotics tested, which may be related to the broader functions of *retS* in regulating a large number of virulence-associated genes in addition to T4P^48^.

Finally, we determined whether the phage-resistance tradeoff relationships with antibiotic sensitivity could be used to predict effective *P. aeruginosa* phage-antibiotic cotreatments. To test this, we performed cotreatment assays combining the three phages (PB1, ABO9, and PhiKZ) with the five antibiotics studied and calculated resistance emergence indexes for each phage-antibiotic pair. Some combinations suppressed resistance emergence, whereas others did not (**Fig. 4d**). Consistent with the *E. coli* results, resistance emergence indexes were significantly negatively correlated with the MICs of phage-resistant strains for both PB1-antibiotic (ρ = 0.95, *P* = 0.01) (**Fig. 4e**), and ABO9–antibiotic cotreatments (ρ = 0.87, *P* = 0.05) (**Fig. 4f**). PhiKZ cotreatments also showed a negative, though weaker, correlation (ρ = 0.71, *P* = 0.18). Together these results suggest that our findings that LPS-driven phage-antibiotic genetic tradeoffs are common and predict resistance suppression that can be generalized from *E. coli* to *P. aeruginosa*.

## Discussion

The global antimicrobial resistance crisis demands rational strategies for new therapies. Although phage therapy has regained attention, its efficacy is limited by the rapid evolution of phage-resistant bacteria. Combining phages with conventional antibiotics enhances treatment efficacy compared to phage monotherapy. However, current combinatorial approaches typically rely on empirical testing of phages capable of infecting the target bacteria alongside selected antibiotics^15,49-51^. Quantitative frameworks like synograms can identify synergistic pairs that enhance initial bacterial killing^19^, yet they fail to address the pressing clinical challenge of preventing the emergence of resistance during treatment. Moreover, these approaches lack mechanistic insight into phage-antibiotic interactions. Here, we establish a mechanistic, predictive framework for the rational design of effective phage–antibiotic combinations grounded in evolutionary tradeoffs. We demonstrate that when phage resistance increases bacterial susceptibility to a specific antibiotic, the corresponding cotreatment not only suppresses bacterial growth but also suppresses resistance emergence. These findings suggests that cotreatments with agents that have evolutionary tradeoffs can be viewed as a form of evolutionary steering, where the strong selection pressure imposed by the phage directs bacterial evolution down a predictable, costly path of losing a cell surface receptor, which then sensitizes the bacterium to a second targeted antibiotic agent that exploits that loss.

The evolutionary tradeoffs we identified are mechanistically predictable because phage resistance often arises through loss-of-function mutations in phage receptor genes. In our sequenced phage-resistant isolates, most carried disruptive mutations in these receptors or regulators of their receptors. LPS, a major component of the Gram-negative bacterial outer membrane, serves as a common receptor for many *E. coli* and *Pseudomonas aeruginosa* phages. We found that mutations disrupting LPS biosynthesis sensitize both *E. coli* and *P. aeruginosa* to specific antibiotics. Consequently, cotreatments combining LPS-targeting phages with these sensitizing antibiotics effectively suppressed the emergence of resistance. Moreover, focusing on LPS-targeting phages provides mechanistic insight into the molecular nature of the tradeoff, as LPS differentially affects the ability of lipophilic molecules to gain cell entry and antibiotic lipophilicity predicted cotreatment outcomes. This mechanistic understanding could allow a clinician to bypass resource-intensive screening and prioritize antibiotic partners based solely on the phage receptor identity and the drug’s biochemical properties. In addition to LPS, which may serve as a broadly applicable target for phage therapy, species-specific tradeoffs are also valuable. Our results demonstrated that in *P. aeruginosa*, resistance to a T4P phage increased susceptibility to piperacillin. The vast majority of *P. aeruginosa* phages target either LPS or T4P, such that our findings suggest that nearly all *P. aeruginosa* phage therapies could be synergistically improved by combining them with colistin/polymyxin B and piperacillin. Finally, while TolC-targeting phages are relatively rare, our findings support the conclusions of a prior study suggesting that resistance to a TolC-targeting phage sensitized *E. coli* to antibiotics expelled by the AcrAB–TolC efflux pump.

While cotreatment outcomes can be predicted by the resistance tradeoffs of receptor mutations, the potential complexity of these mutations should not be overlooked. For example, seven of the eight *P. aeruginosa* clones resistant to LPS-targeting phages carried large genomic deletions spanning not only the LPS-related gene (*galU*) but also the adjacent efflux pump genes (*mexX* and *mexY*). These co-occurring deletions conferred the expected sensitivity to polymyxin B and colistin (mediated by *ΔgalU*) while also generating a novel sensitivity to amikacin (mediated by *ΔmexXY*). This finding suggests that enhanced antibiotic susceptibility can be achieved through the co-loss of neighboring genes of the receptor gene. Furthermore, the identification of a regulatory mutation (*ΔretS*) in a PhiKZ-resistant *P. aeruginosa* strain, which regulates the GacS/GacA two-component system^48^, introduces the complexity of tradeoffs caused by regulatory mutations that could have more pleiotropic effects. Finally, the existence of phages that exhibit dual receptor dependency, such as those relying on both LPS and TolC (e.g. U136B)^22^, can lead to competing tradeoffs with a single antibiotic (e.g., tetracycline), potentially complicating cotreatment outcomes. Thus, there are scenarios where the complexity associated with phage resistance can be exploited to achieve additional tradeoffs.

The significant correlation between the genetic tradeoff (increased antibiotic sensitivity following receptor loss) and the resistance emergence index confirms that this framework is a reliable and portable predictor of treatment success across both *E. coli* and *P. aeruginosa*. This constitutes a substantial advance over efficacy-focused models, which did not significantly correlate with resistance emergence. Critically, our rational design framework suggests that these phage-antibiotic combinations create a “double threat” for the pathogen, offering robust clearance potential. First, the phage decimates the wild-type population. Second, the resistant mutants modify their primary permeability barrier, rendering them hypersensitive to antibiotics. In the case of LPS-targeting mutants, there is potential for a third threat because LPS truncation is well-established to severely attenuate virulence and sensitize Gram-negative bacteria to host innate immunity (e.g., complement-mediated lysis and phagocytosis)^52,53^. Thus, bacteria escaping the phage are immediately compromised against both the antibiotic and the host immune system.

To translate this strategy into clinical reality, future work must focus on platform generalization to other priority Gram-negative pathogens like *Acinetobacter baumannii* and *Klebsiella pneumoniae*, where LPS is also a dominant phage receptor^54-56^. Furthermore, pharmacokinetic and pharmacodynamic (PK/PD) studies are essential to validate these combinations *in vivo*, ensuring that the antibiotic dosage reliably achieves the necessary concentration at the site of infection to eliminate the newly sensitized mutant population without inducing patient toxicity. Other physiological considerations can add further complexity. For example, the gut environment can change bacterial gene expression, thus changing both phage and antibiotic susceptibility^57^. Future work of phage-antibiotic cotreatment test *in vivo* will thus be essential to assess their robustness in realistic infection contexts. By prioritizing the evolutionary path dictated by the phage receptor, our approach provides the mechanistic basis for the precise and rational design of combination therapies, moving phage therapy closer to its full potential as a targeted anti-resistance tool.

## Materials and methods are detailed in the supplementary information

